# pH dependence of C•A, G•A and A•A mismatches in the stem of precursor microRNA-31

**DOI:** 10.1101/2021.10.25.465784

**Authors:** Anita Kotar, Sicong Ma, Sarah C. Keane

**Affiliations:** Biophysics Program, University of Michigan, 930 N. University Avenue, Ann Arbor, MI 48109, USA; Department of Chemistry, University of Michigan, 930 N. University Avenue, Ann Arbor, MI 48109, USA; Current Address: Slovenian NMR Centre, National Institute of Chemistry, Hajdrihova 19, SI-1000 Ljubljana, Slovenia

**Keywords:** microRNA, helical stem, base pair mismatch, solution NMR spectroscopy

## Abstract

MicroRNAs (miRNAs) are important regulators of post-transcriptional gene expression. Mature miRNAs are generated from longer transcripts (primary, pri- and precursor, pre-miRNAs) through a series of highly coordinated enzymatic processing steps. The sequence and structure of these pri- and pre-miRNAs play important roles in controlling their processing. Both pri- and pre-miRNAs adopt hairpin structures with imperfect base pairing in the helical stem. Here, we investigated the role of three base pair mismatches (A∙A, G∙A, and C∙A) present in pre-miRNA-31. Using a combination of NMR spectroscopy and thermal denaturation, we found that the three base pair mismatches displayed unique structural properties, including varying dynamics and sensitivity to solution pH. These studies deepen our understanding of how the physical and chemical properties of base pair mismatches influence RNA structural stability.

**Graphical abstract:** 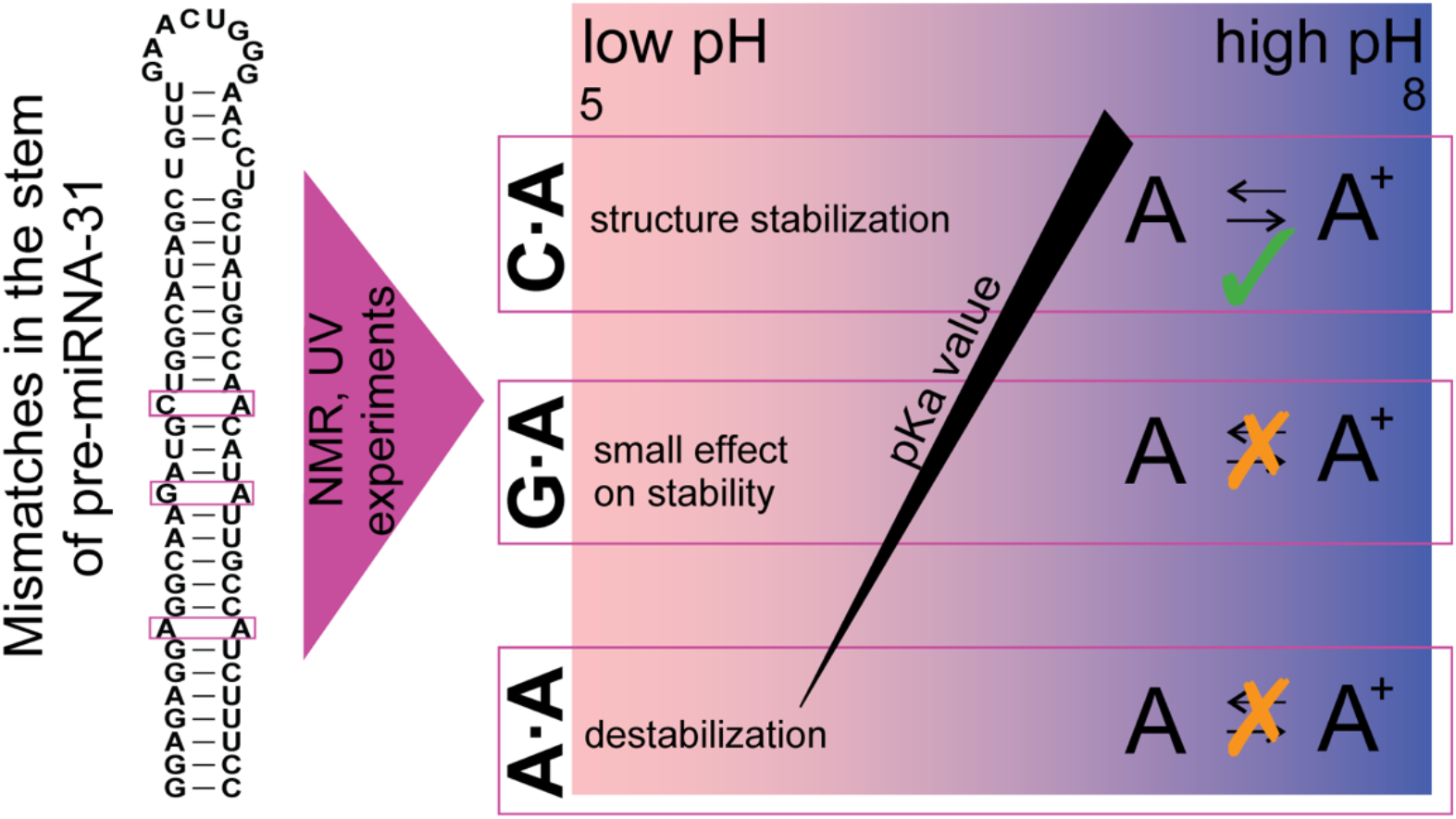

## 1. Introduction

MicroRNAs (miRNAs) are short (~22 nucleotide) non-coding RNAs that play an important role in the regulation of gene expression through targeting messenger (m) RNAs for post-transcriptional gene silencing [1]. Post-transcriptional regulation of miRNA biogenesis is a multistep process that starts when a long primary (pri-) miRNA is enzymatically processed by Drosha/DGCR8 to generate the precursor (pre-) miRNA. The pre-miRNA is exported from the nucleus where it is further processed by the cytoplasmic Dicer enzyme into the mature miRNA [1]. One strand of the mature miRNA duplex is loaded into the RNA-induced silencing complex (RISC), where it serves as template for complementary recognition of the target mRNA. It has been suggested that one third of genes in the genome are regulated by miRNAs [2–4], impacting diverse biological processes [5–7]. miRNA processing is tightly regulated to ensure accurate gene expression. The sequence and structure of both pri- and pre-miRNAs play important roles in regulation of miRNA processing as both pri- and pre-miRNAs are differently recognized by the processing enzymes [8–13]. The presence of stable basal stems in pri-miRNAs and flexible apical loops in pri-/pre-miRNAs lead to more efficient processing by the biogenesis machinery [14]. The effects of other structural elements like mismatches within stems of pri-/pre-miRNAs on processing are not as well characterized, but are emerging as important regulators of miRNA maturation [15]. This is especially important given that more than two-thirds of human pre-miRNAs contain at least one base pair mismatch (**Fig. 1**).

**Figure 1.**
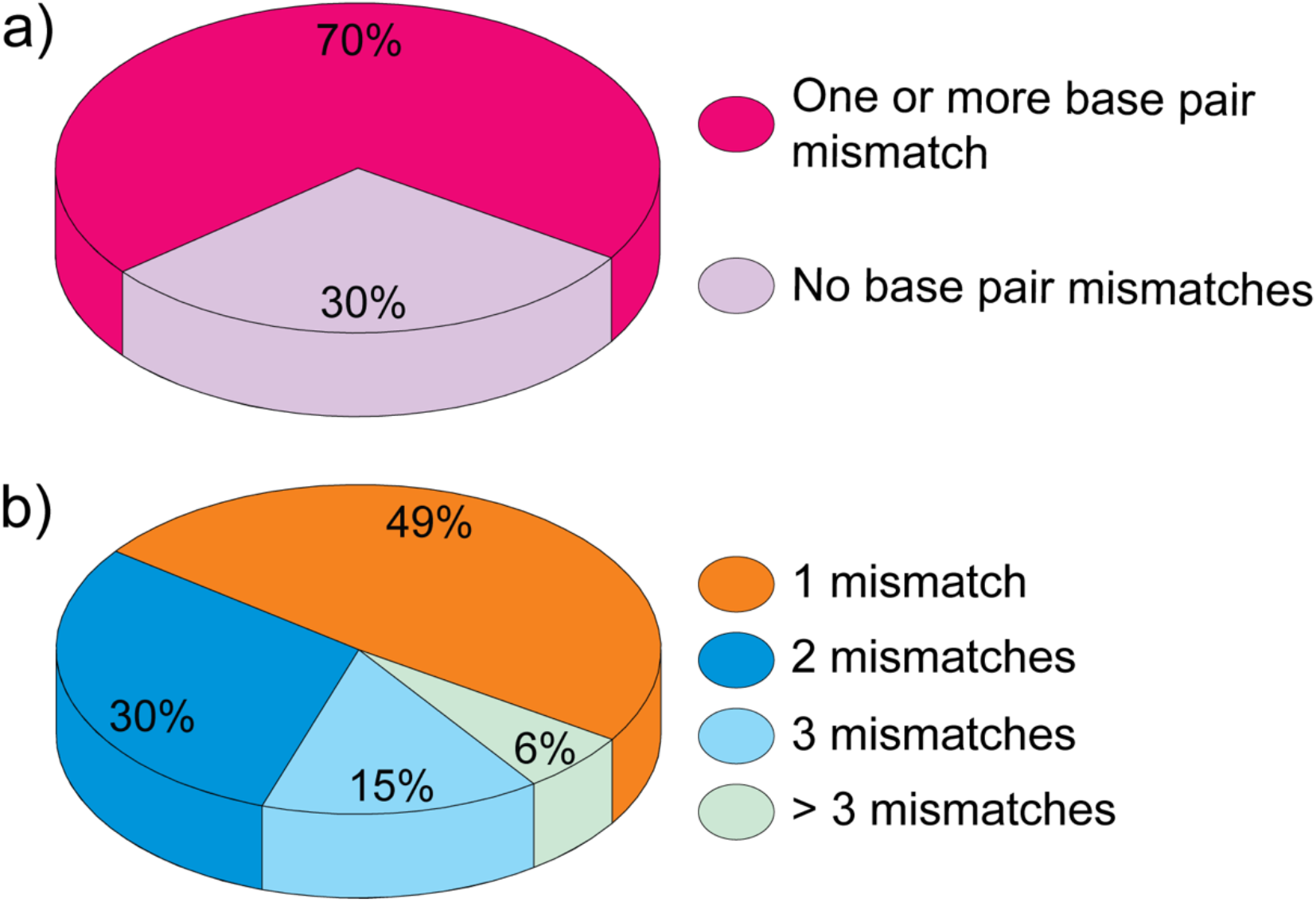
Statistics of base pair mismatches in human pre-miRNA stems. Fraction of RNAs a) with and without mismatches and b) breakdown of the number of mismatches present in pre-miRNA stems.

Mismatches and wobble base pairs in the upper stem of pri-miRNAs can impact the efficiency and accuracy of miRNA processing [16]. Importantly, some non-canonical base pair geometries in pri-/pre-miRNAs can be induced by altering solution conditions, such as pH. An A^+^-G mismatch was found in the stem-loop region of pre-miRNA-21 exited states, which enhanced Dicer processing over its ground conformational state [17]. While in single stranded regions of RNA, the nucleobases are typically uncharged, with reported pK_a_ values of 3.5, 4.2, and 9.2 for A, C and G/U, respectively [18]. In higher order structural environments like mismatches, the pK_a_ values can be shifted towards neutrality which leads nucleobases to adopt uncommon protonated states that can facilitate different non-canonical interactions [19].

We examined the base pair mismatches in the stem of the evolutionarily conserved pre-miRNA-31, which is involved in maintaining fertility, embryonic development, bone formation, and myogenesis [20]. miRNA-31 is known to interact with a series of target genes and pathways; thus its mis-regulation has been connected to various diseases like cancer, autoimmune diseases, and heart conditions such as atrial fibrillation [20–24]. Notably, in tumorigenesis miRNA-31 can act as an enhancer of tumor development and progression (in lung, colorectal, non-small-cell lung, head and neck squamous cell, and esophageal squamous cell cancers) as well as a tumor suppressor (in breast, ovarian, and prostate cancers and in hepatocellular and gastric carcinoma) [23]. Additionally, miRNA-31 regulates diverse processes during embryonic implantation and development as well as promotes early sperm development [20]. Moreover, miRNA-31 can positively regulate the proliferation, differentiation and cell activity of keratinocytes, which are important functions connected to various skin diseases as well as wound healing [22, 24]. Pre-miRNA-31 contains three base pair mismatches (C∙A, G∙A, and A∙A) in its helical stem. Using NMR and UV spectroscopic methods, we evaluated the structural characteristics of these base pair mismatches and analyzed how their properties change when the pH of the buffering solution was altered.

## 2. Materials and methods

### 2.1 Cataloging of mismatches in human pre-miRNAs

All 1,917 pre-miRNA sequences were downloaded from the miRBase human microRNA database [25] and secondary structures were predicted using the RNAstructure web server [26] with default parameters. We defined the stem as regions between the 5ʹ-most base paired nucleotides (distal from the apical loop) and the last set of base paired nucleotides which are closest to the Dicer cleavage site (proximal to the apical loop). Almost all (1,883/1,917) of the extracted pre-miRNA structural information was based on the predicted lowest energy structure. However, in a few cases (34/1,917) the data was based on the lowest energy structure which maintained a canonical stem-loop structure. All relevant information, including the identity of base pairs (A-U, G-C, G-U), single nucleotide mismatches (C∙A, A∙G, A∙A, C∙C, U∙C, U∙U, G∙G), and unpaired nucleotides (including internal loops and single nucleotide bulges), was recorded and analyzed.

### 2.2 Construct design and template preparation

RNAs examined in this study are listed in **Table 1**. DNA oligonucleotides were purchased from Integrated DNA Technologies. The DNA templates were ordered with 2ʹ-*O*-methoxy modifications at the two 5ʹ-most positions to reduce non-templated transcription [27]. The DNA templates for *in vitro* transcription were created by annealing the DNA oligonucleotides (BottomA: 5ʹ-mGmGAAAGATGGCAATCTCTTGCCTCCTCTCC*<u>TATAGTGAGTCGTATTA</u>*-3ʹ, BottomB: 5ʹ-mGmGCAATATGTTGGTCTCCCAGCATCTTGCC*<u>TATAGTGAGTCGTATTA</u>*-3ʹ; where m denotes 2ʹ-*O*-Me modification of the oligonucleotide and italicized nucleotides correspond to the sequence complementary to the T7 promoter) with an oligonucleotide corresponding to the T7 promoter sequence (5ʹ-TAATACGACTCACTATA-3ʹ). Templates were prepared by mixing the desired DNA oligonucleotide (40 μL, 200 μM) with the complementary oligonucleotide to T7 promoter sequence (20 μL, 600 μM) together, boiling for 3 min, and then slowly cooling to room temperature. The annealed template was diluted with H_2_O prior to use to produce the partially double-stranded DNA templates at a final concentration approximately 8 μM.

**Table 1.**
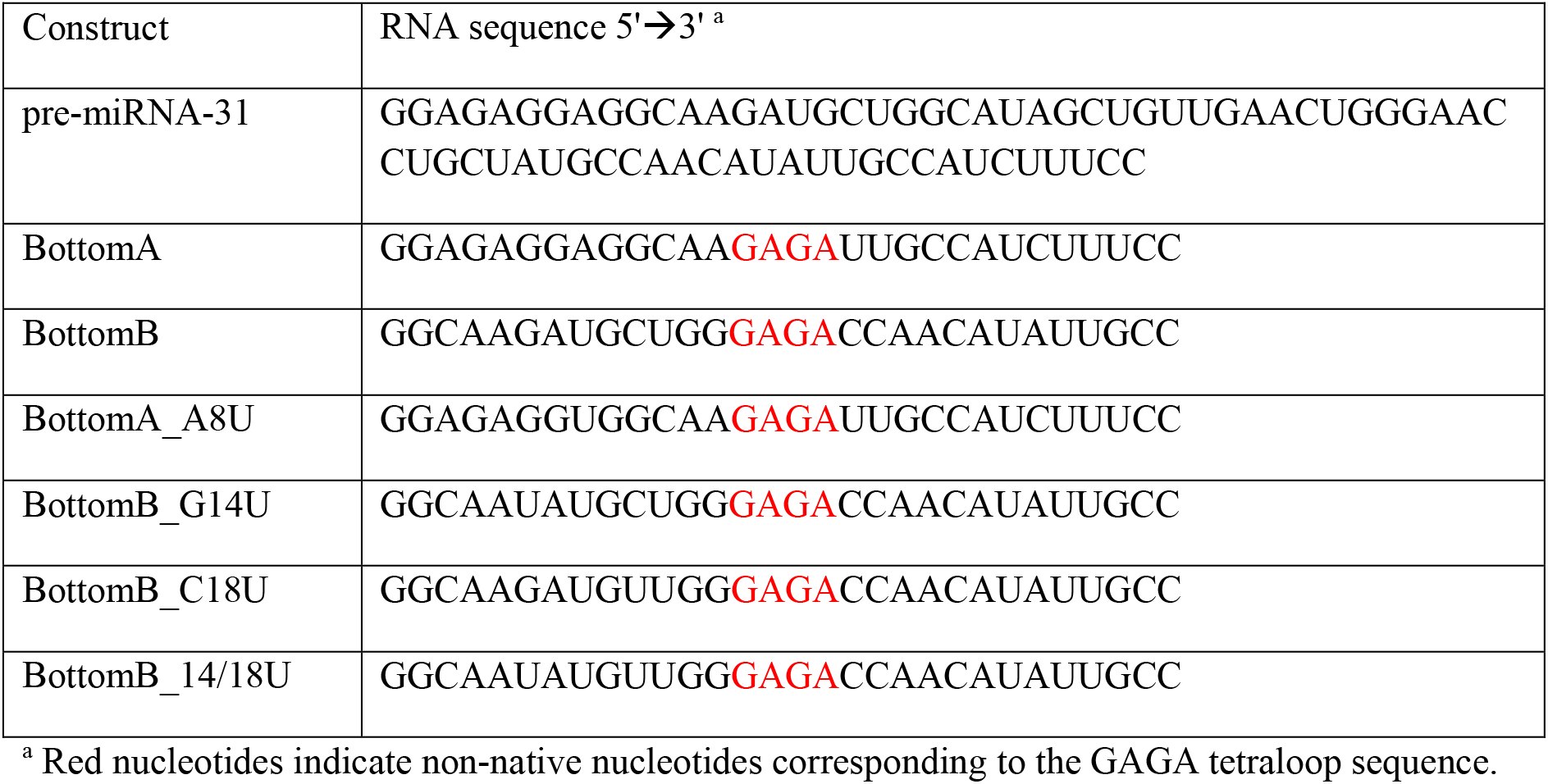
RNA constructs.

### 2.3 RNA Preparation

pre-miRNA-31, BottomA and BottomB RNAs (**Table 1**) were prepared by *in vitro* transcription in 1X transcription buffer [40 mM Tris base, 5 mM DTT, 1 mM spermidine, and 0.01% Triton-X (pH=8.5)] with addition of 3−6 mM NTPs, 10−20 mM MgCl_2_, 30−40 ng/μL DNA template, 0.2 unit/mL yeast inorganic pyrophosphatase (New England Biolabs) [28], ∼15 μM T7 RNA polymerase, and 10−20% (v/v) DMSO. Reaction mixtures were incubated at 37 °C for 3−4 h, with shaking at 70 rpm, and then quenched using a solution of 7 M urea and 500 mM EDTA (pH=8.5). Reactions were boiled for 3 min and then snap cooled in ice water for 3 min. The transcription mixture was loaded onto preparative-scale 14% (pre-miR-31) or 18% (BottomA, BottomB) denaturing polyacrylamide gels for purification. Gel slices containing the target RNA were crushed and soaked in TBE buffer for 24-48 h to extract the RNA. Eluted RNA was filtered and spin concentrated, washed with 2 M high-purity sodium chloride, and exchanged into water using Amicon-15 Centrifugal Filter Units (Millipore, Sigma). RNA quality was verified by running the purified RNA on an analytical 14% (pre-miR-31) or 18% (BottomA, BottomB) denaturing polyacrylamide gel.

The modified RNAs with stabilizing mutations (BottomA_A8U, BottomB_G14U, BottomB_C18U and BottomB_14/18U) and site-specific ^13^C- or ^15^N-labeled RNAs were chemically synthesized using DMT-on phosphoramidite chemistry on K&A Laborgeraete GbR DNA/RNA Synthesizer H-8. Samples were purified using the Glen-Pak RNA Cartridge Purification (DMT-ON) protocol [29]. RNAs were washed with 2 M high-purity sodium chloride, and exchanged into water using Amicon-15 Centrifugal Filter Units (Millipore, Sigma).

Purified and desalted RNAs were refolded by heating in boiling water for 3 min, followed by incubation on ice for 3 min. For NMR experiments in 100% D_2_O, the RNA samples were lyophilized and dissolved in 100% D_2_O (99.8%, Cambridge Isotope Laboratories, Inc.) and transferred to an Eppendorf tube containing lyophilized buffer (50 mM K-phosphate buffer (pH=7.5 or pH=5.8) and 1 mM MgCl_2_) with RNA concentration of 0.4 mM. Other NMR samples were prepared in 90%/10% H_2_O/D_2_O with 50 mM K-phosphate buffer (pH=7.5 or pH=5.8), 1 mM MgCl_2_, and 0.2-0.5 mM RNA concentrations.

### 2.4 NMR experiments

2D ^1^H-^1^H NOESY, ^1^H-^1^H TOCSY, and ^1^H-^13^C HMQC spectra for NMR assignment were recorded at 30 °C for BottomA, and at 37 °C for BottomB. The following experimental parameters were used for the ^1^H-^1^H NOESY: pulse sequence noesyphpr [30], ds=16, ns=32-128, sw(F2)=10.0138-10.7724, sw(F1)=10.0138-10.7724, TD(F2)=8129, TD(F1)=68-800, O1=4.708, D1=2-5 s, *τ*_m_=400-500 ms; the ^1^H-^1^H TOCSY: pulse sequence mlevgpph19 [31–33], ds=32, ns=128-256, sw(F2)=10.0138-10.2427, sw(F1)=10.0138-10.2427, TD(F2)=2048, TD(F1)=256-360, O1=4.708, D1=2 s, *τ*_m_=80 ms; and the ^1^H-^13^C HMQC: pulse sequence hmqcphpr [34], ds=16, ns=304, sw(F2)=8.778, sw(F1)=50.0, TD(F2)=1058, TD(F1)=80, O1=4.705, O2=148.00, D1=1.5 s. NMR spectra were collected on 600 and 800 MHz Bruker AVANCE NEO spectrometers equipped with a 5 mm TCI cryogenic probe and on a 600 MHz Bruker AVANCE DRX NMR spectrometer equipped with a 5 mm 5mm PFG cryoprobe (University of Michigan BioNMR Core). NMR data were processed with NMRFx [35] analyzed with NMRViewJ [36] and MestReNova 12.0.0-20080 [37]. ^1^H chemical shifts were referenced to water and ^13^C chemical shifts were indirectly referenced from the ^1^H chemical shift [38].

### 2.5 pH titrations and pKa measurements

BottomA and BottomB samples for the pH titration NMR experiments were prepared with 0.2 mM concentration of RNAs in 10% D_2_O, and 1 mM MgCl_2_. Initial pH values were set to 7.5 with 100 mM NaOH using SevenEasy Mettler Toledo pH meter. For the pH titration series, the samples were titrated with 50 mM HCl and after each addition of acid the pH was checked using the pH meter. All solutions were freshly prepared the same day NMR spectra were acquired.

The changes in ^1^H chemical shifts were followed during the pH titration. The graphs of chemical shift changes versus pH values were used to determine pKa values using **Equation 1** [39]:

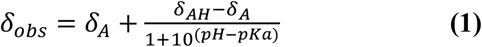

where δ_A_ is the chemical shift at high pH, δ_AH_ is the chemical shift at low pH, and δ_obs_ is the observed chemical shift at a given pH.

### 2.6 Thermal denaturation of RNA and data analysis

UV-thermal denaturation experiments were performed using an Agilent Cary UV-Vis Multicell Peltier spectrometer with a heating rate of 0.5 °C per min between 10 to 95 °C and 95 to 10 °C. Data points were collected every 1 °C with absorbance detection at 260, 295 and 330 nm. RNA samples (20 μM) were prepared in 50 mM K-phosphate buffer (pH=7.5 or pH=5.8). The melting profiles (at 260 nm) revealed reversible single-transition unfolding and were analyzed using a two-state model with sloping baselines [40] (**Equation 2**):

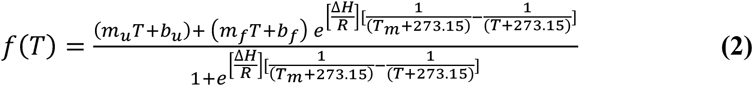

where m_u_ and m_f_ are the slopes of the lower (unfolded) and upper (folded) baselines, b_u_ and b_f_ are the y-intercepts of the lower and upper baselines, respectively. ΔH (in kcal/mol) is the enthalpy of unfolding and T_m_ (in °C) is the melting temperature, R is the gas constant (0.001987 kcal/(Kmol)). Experiments were performed in duplicate and were in good agreement.

## 3. Results

### 3.1 70% of human pre-miRNAs contain at least one base pair mismatch in the stem

We predicted the secondary structures of all pre-miRNAs using the RNAstructure web server [26] and analyzed the occurrence of base pair mismatches. We found that approximately 70% (1,351/1,917) of human pre-miRNAs contain at least one base pair mismatch in the stem (**Fig. 1a**). Most pre-miRNAs were predicted to contain a single base pair mismatch (49 %), followed by those predicted to contain two (30%). Pre-miRNAs containing three base pair mismatches accounted for 15% of those surveyed, while relatively few (6%) of pre-miRNAs were predicted to contain more than three base pair mismatches (**Fig. 1b**). C∙A mismatches are found in 41% (559/1,351) of pre-miRNAs containing at least one mismatch. A summary of the presence of other mismatches is as follows: U∙C mismatches (29%, 396/1,351); G∙G mismatches (23%, 307/1,351); U∙U mismatches (22%, 302/1,351); G∙A mismatches (18%, 247/1,351); A∙A mismatches (13%, 174/1,351), and C∙C mismatches (11%, 142/1,351). Pre-miRNA-31 is predicted to contain three mismatches, C∙A, G∙A and A∙A, in its stem region (**Fig. 2a**).

**Figure 2.**
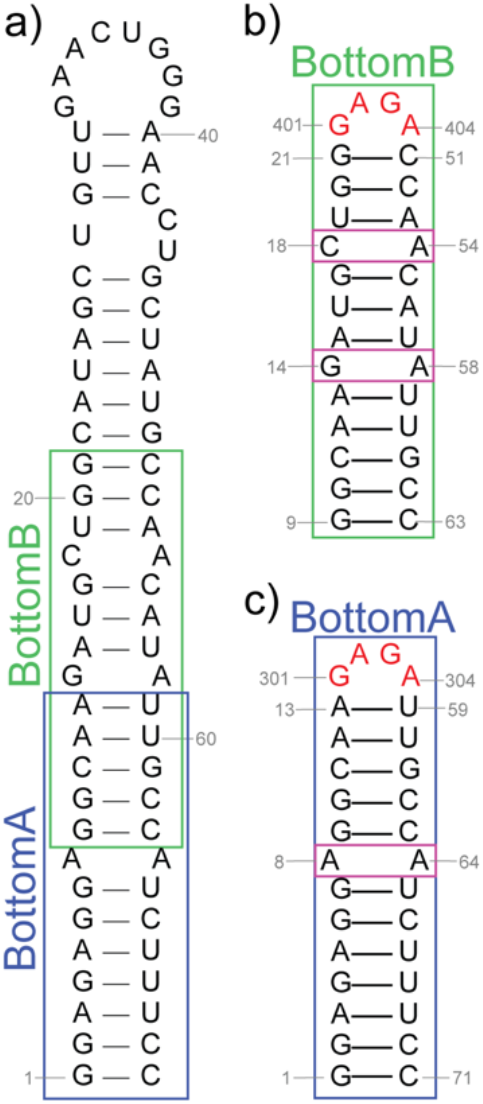
Predicted secondary structure of pre-miRNA-31. Secondary structure of a) pre-miR-31, b) BottomB, and c) BottomA RNA constructs. Colored rectangles denote the location of BottomA and BottomB constructs relative to pre-miRNA-31. The mismatches of interest are boxed with magenta. The secondary structures were predicted using the RNAStructure webserver [26] and were rendered using RNA2Drawer [77].

### 3.2 NMR data reveal the pre-miRNA-31 base pair mismatches have different conformations

In the predicted pre-miRNA-31 secondary structure (**Fig. 2a**), we focused on the helical stem region containing three base pair mismatches. We designed two RNAs, BottomA and BottomB, to examine the structure and properties of these base pair mismatches. The stem of BottomA is composed of four AU, six GC base pairs, and two GU base pairs. BottomA also contains one A∙A mismatch. The stem of BottomB consists of five AU and six GC base pairs. BottomB also has one C∙A and one G∙A mismatch. Both BottomA and BottomB RNAs contain a non-native GAGA tetraloop designed to cap the helical stem (**Fig. 2b,c**). The GAGA-tetraloop is structurally stable and yields unambiguous and characteristic patterns of signals in 2D ^1^H-^1^H NOESY spectra [41–42], which facilitated resonance assignments. The observed NOE signals of BottomA and BottomB revealed that both RNAs adopted an A-form helical stem with a properly folded GAGA tetraloop (**Fig. 3**). Both RNAs also adopted the same conformations as observed in full length pre-miRNA-31 as is evidenced by the matching signals in NOESY spectra of both BottomA and BottomB when compared to the full-length pre-miRNA-31 (**Fig. S1**).

**Figure 3.**
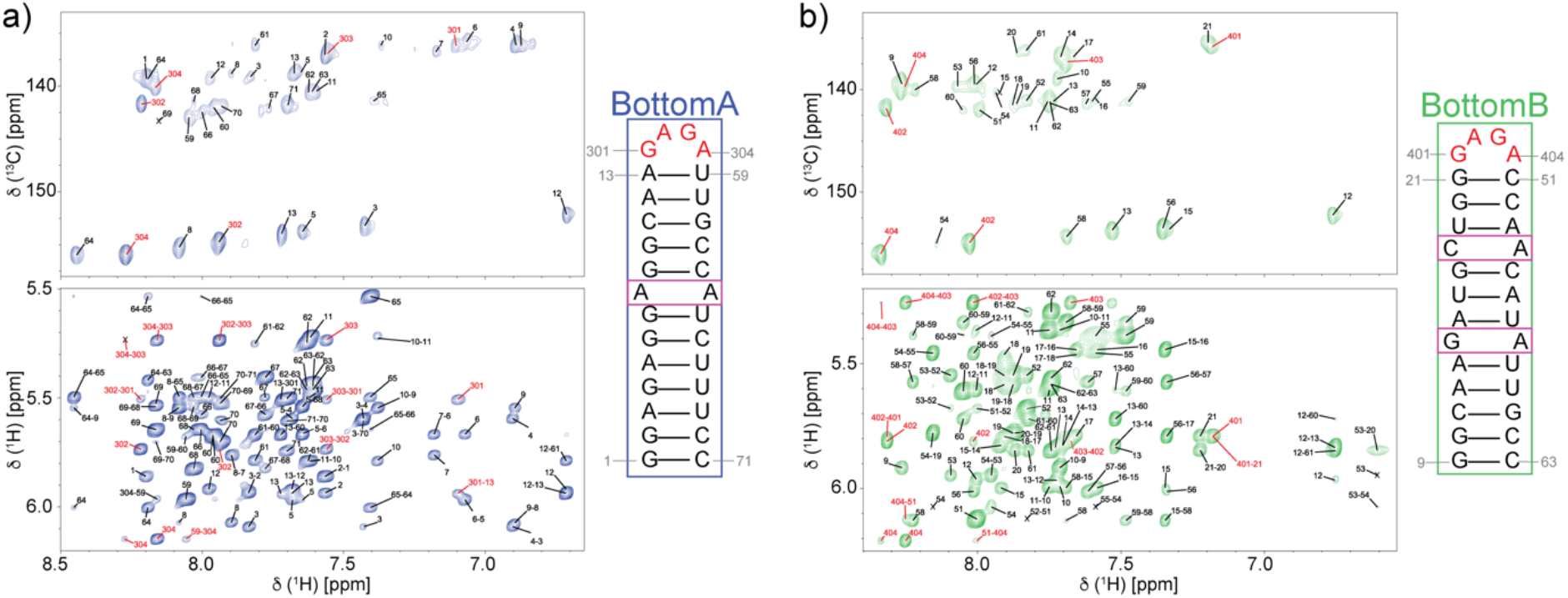
Assigned chemical shifts of BottomA and BottomB RNAs. ^1^H-^13^C HMQC (top) and ^1^H-^1^H NOESY (bottom) spectra of a) BottomA and b) BottomB RNAs. The secondary structure of each oligo is shown to the right of the spectra. The signals assigned to GAGA tetraloop are colored red. NMR spectra were recorded at 0.4 mM RNA concentration, 50 mM K-phosphate buffer, pH=7.5, 1 mM MgCl_2_ and 100% D_2_O.

The signals of nonexchangeable protons of BottomA and BottomB were assigned based on analysis of 2D ^1^H-^1^H NOESY, 2D ^1^H-^1^H TOCSY, and ^1^H-^13^C HMQC spectra (**Fig. S2**). 2D ^1^H-^1^H NOESY spectra were used to assign through space correlations and to obtain sequential connectivities in the A-helical stem regions of BottomA and BottomB. The characteristic NOEs of GAGA tetraloop residues were used as reference points which helped to facilitate the resonance assignments of BottomA and BottomB via the established principles of a sequential walk (**Fig. 3**). The NOE signals from C2 protons of adenosine residues were crucial in identifying cross-strand NOE connectivities and consequently identifying mismatch conformations. In the case of the A8∙A64 mismatch in BottomA, we observed cross-strand connectivities between A8.H2 and 65.H6 as well as A64.H2 and G9.H8. We also observed strong sequential NOE connectivities with neighboring residues for both A8 and A64. These results indicate that A8 and A64 are oriented inside the stem, stacked relative to their neighboring residues.

Similarly, for the G14∙A58 mismatch in BottomB we observe NOE cross-strands connectivities between A58.H2 and A15.H2. Interestingly, for G14 we observe clear sequential NOE connectivities with A15, while the sequential NOE connectivities between G14 and A13 could not be confirmed due to either missing signals or signal overlap in the aromatic-anomeric region. Taken together, these findings indicate that G14 is not only oriented inside the stem and engaged in common A-form RNA staking interactions with sequential neighbors but also can adopt conformations that disturb the ideal staking arrangements or can position itself outside the stem. For the C18∙A54 mismatch, we observed cross-strand NOE signals between A54.H2 and U19.H1’. We noted that signals corresponding to C18 as well as neighboring A53 and C55 nucleotides exhibit very broad cross-peaks in aromatic-aromatic and aromatic-anomeric regions of NOESY spectrum. We did not detect similar broadening for proton signals of nucleotides in other two mismatches or for the other protons in BottomA and BottomB. These observations indicate chemical exchange between different structural conformations happening in the region of the C18∙A54 mismatch.

### 3.3 Stabilization of the structure in the C·A mismatch at lower pH

Protonation of nucleobases can promote formation of non-canonical base pairs in mismatches and other structural elements within RNA molecules and has been connected to many important biological functions [19]. Indeed, there are several examples showing that protonated adenosines may play important mechanistic roles in adenosine deamination, regulation of translational recording, ligand binding in RNA aptamers and riboswitches, and processing of microRNAs, among others [17, 43–46]. To explore in more detail how pH influences the structures and the dynamic behaviors of mismatches in pre-miRNA-31, we analyzed 1D ^1^H NMR spectra of BottomA and BottomB collected at different pH values (**Fig. 4**). During the stepwise titration of BottomA with HCl, from pH=7.5 to pH=5.2, we did not observe significant changes in chemical shifts in the imino region, only slight broadening as the pH was lowered (**Fig. 4a**). In BottomA, we were interested in characterizing the pH-dependent changes of residues in the A8∙A64 mismatch. Unfortunately, A8.H8 and A8.H2 signals were too overlapped for a conclusive analysis, however, the A64.H2 proton signal was nicely resolved at 8.41 ppm (pH=7.5). The chemical shift of the A64.H2 signal changes to 8.39 ppm at lower pH values and becomes slightly broader. These results were confirmed in the ^1^H-^15^N HSQC spectra where the A64.H2-A64.N1 cross-peak is much broader at pH=5.8 compared to pH=7.5 (**Fig. S3**). At 10.5 ppm we detected a signal from G301.H1 (partially overlapped with the signal of the GU base pair) which forms a G∙A base pair within the GAGA tetraloop. The G301.H1 signal became sharper at lower pH. This is consistent with previous observations and indicates that the G301.H1 proton exchange with solvent is even slower at low pH [41, 47].

**Figure 4.**
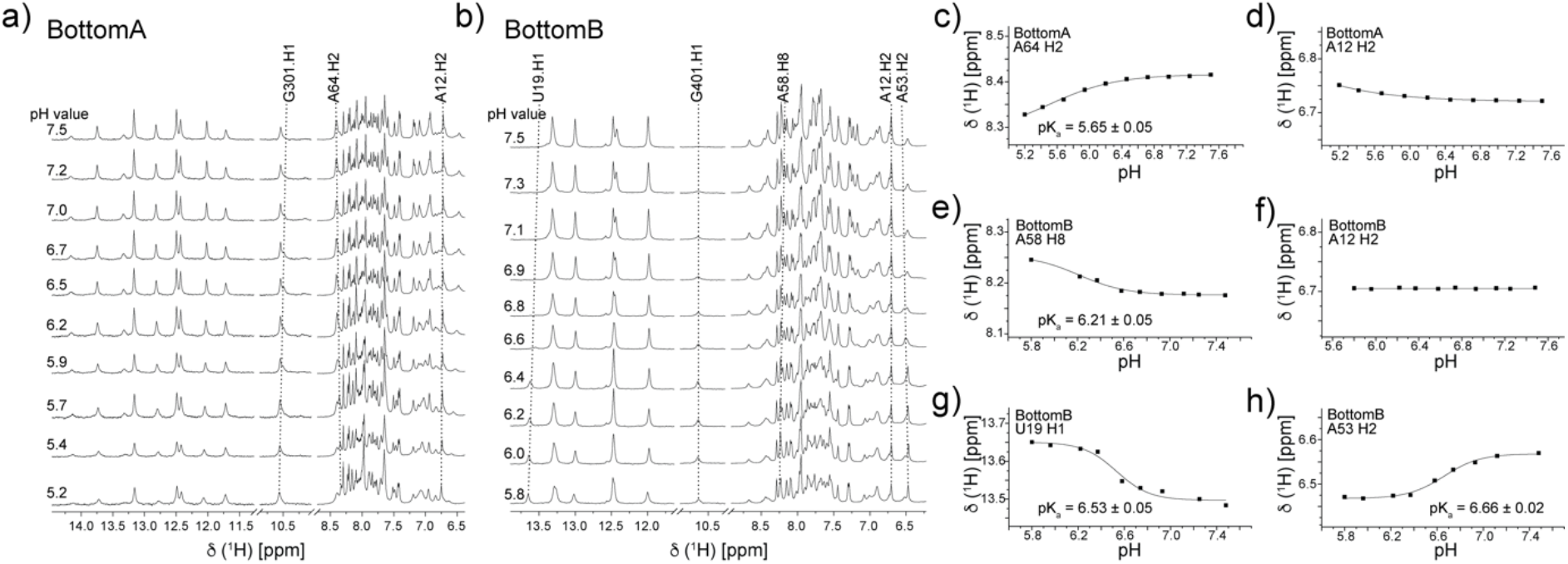
pH dependence of chemical shifts. The imino and aromatic regions of ^1^H NMR spectra of a) BottomA and b) BottomB RNAs at different pH. The pH values are indicated on the left side of the spectra. The assignments of selected signals are shown above the spectra and the dotted lines follow the changes in the chemical shifts. Effect of pH on chemical shift for c) BottomA A64 H2, d) BottomA A12 H2, e) BottomB A58 H8, f) BottomB A12 H2, g) BottomB U19 H1 and h) BottomB A53 H2. pK_a_ values derived from these titrations are indicated within each panel. NMR spectra were recorded at 0.2 mM of RNAs, 1 mM MgCl_2_, 90%/10% H_2_O/D_2_O at 37 °C.

Analysis of the pH titration of BottomB revealed more significant changes in both the imino and aromatic regions of 1D ^1^H NMR spectra compared to BottomA (**Fig. 4b**). Consistent with our observations in BottomA, the chemical shift of the G401.H1 signal is at 10.64 ppm, which indicates the stabilization of the G∙A base pair from the GAGA tetraloop. The most pronounced change in the NMR spectra during the titration was a sharpening of a signal at 13.65 ppm (pH=5.8) assigned to U19.H1. This signal was very broad at higher pH (barely detectable above the baseline) and became narrow and well-resolved at lower pH. We observed a comparable effect for the A53.H2 signal at 6.47 ppm (pH=5.8) which became sharp and well-resolved as the pH was lowered. The signals assigned to C18 from the C∙A mismatch were too overlapped to analyze their pH dependence. Interestingly, in the NOESY spectrum recorded at pH=5.8 (**Fig. S4**), cross-peaks for the nucleotides in the C∙A mismatch and the neighboring residues were more narrow and well-defined compared to their broad cross-peaks observed at pH=7.5 (**Fig. 3b**). This indicates that at lower pH the chemical exchange between different conformations around the C∙A mismatch is not present and a single conformation is stabilized.

For the G14∙A58 mismatch, we observed a broadening of the A58.H8 signal when the pH was lowered. Interestingly, in ^1^H-^15^N HSQC spectra we observed much broader H2-N1 cross-peaks even at pH=7.5 for A54 and A58 compared to A64 (**Fig. S5**). We prepared a site-specifically labeled BottomB RNA with a ^15^N1 label at A54. In this sample (pH=7.5), we observed two broad cross-peaks in the ^1^H-^15^N HSQC spectrum, one with a ^15^N chemical shift of 235.4 ppm, expected for an adenine N1 atom, and the other with a ^15^N chemical shift of 156.6 ppm, which is characteristic of protonated N1 atoms (**Fig. 5a**). When the pH was lowered to 5.8, only a single ^1^H-^15^N HSQC cross-peak (δ ^15^N=156.6 ppm) was observed (**Fig. 5b**). The A54.N1-A54.H1 cross-peak at low pH is narrower and more defined in both the ^1^H and ^15^N dimensions compared to the cross-peak at pH=7.5. Additionally, comparison of the ^1^H-^13^C HSQC spectra of BottomB at low and high pH revealed an upfield shift of A54.C2 from 155 ppm (pH=7.5) to 147 ppm (pH=5.8) (**Fig. 5c**). We did not observe similar changes in chemical shifts for other C2 atoms, indicating that only A54 is protonated at low pH. However, the A58 H2-C2 cross-peak is broader at pH=5.8 compared to pH=7.5 (**Fig. 5c**). It is also important to note that also the H2-N1 cross-peak in the ^1^H-^15^N HSQC spectrum of A58 was too broad to detect at pH=5.8 which indicates that A58 is engaged in intermediate chemical exchange between its protonated and non-protonated form. Furthermore, our NMR data show that the sharpening of the A53.H2 signal at lower pH is due to the dynamic behavior in the C∙A mismatch rather than protonation of A53, since the ^13^C chemical shift of the A53 H2-C2 cross-peak is 151.7 ppm, consistent with a non-protonated adenine residue (**Fig. 5c**). Moreover, as we can clearly observe from ^1^H-^13^C HSQC spectra recorded at pH values of 7.5 and 5.8, the C8 ^13^C and H8 ^1^H chemical shifts of A53 (Δδ^13^C 0.1 ppm, Δδ^1^H 0.03 ppm) are resistant to changes in pH compared to those for A54 where we observe a clear pH dependence (Δδ^13^C 1.9 ppm, Δδ^1^H 0.25 ppm) (**Fig. S6**). Collectively, these results strongly suggest that a population of A54 is already protonated at pH=7.5 and virtually completely protonated at pH=5.8 (**Fig. 5**, **S6**).

**Figure 5.**
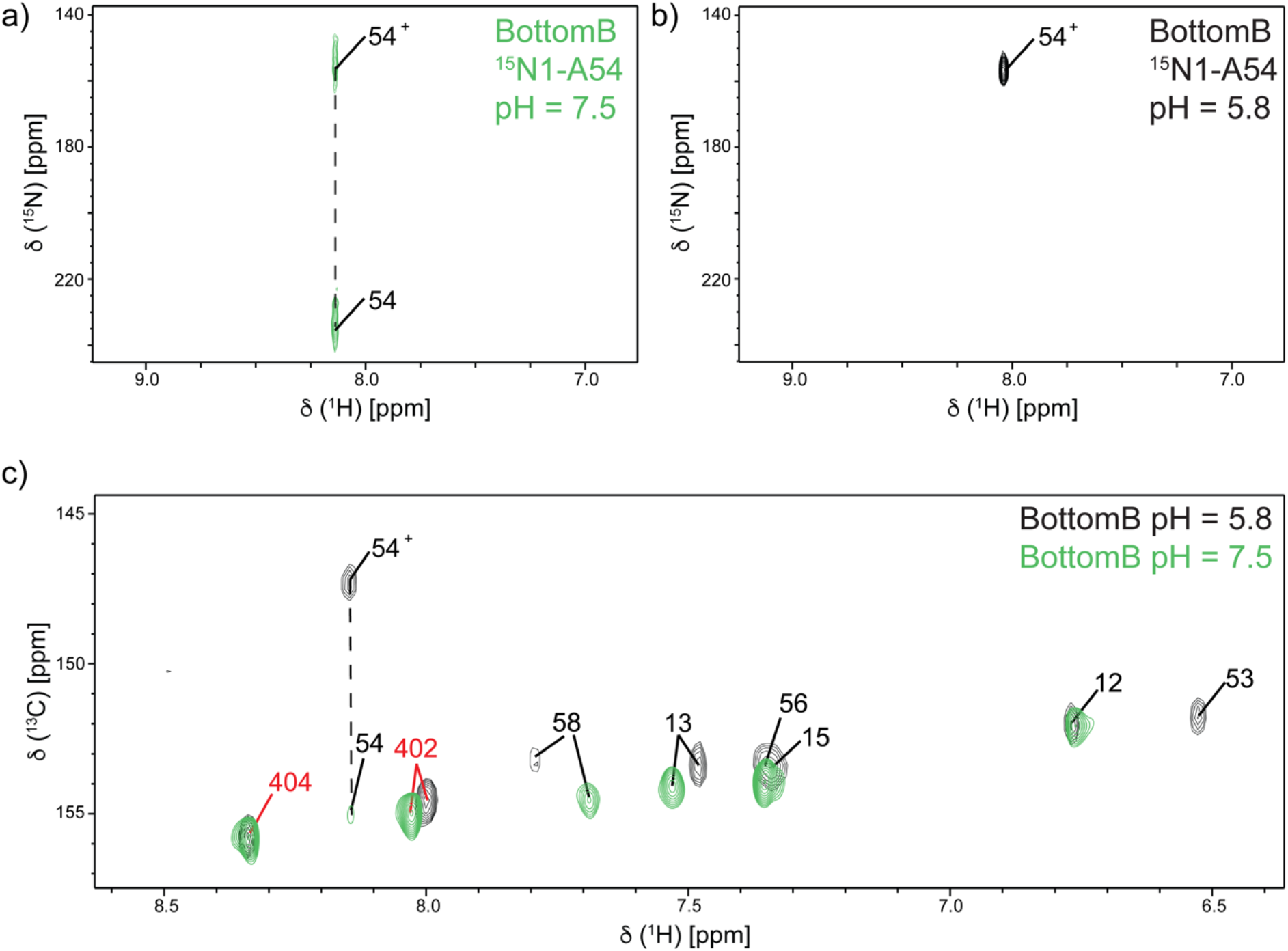
Protonation of A54 is favored at low pH. H2-N1 cross-peak of 100% ^15^N1 labeled A54 in ^1^H-^15^N HSQC spectra recorded at a) pH=7.5 and b) pH=5.8. H2-C2 cross-peaks of BottomB in ^1^H-^13^C HSQC spectra c) at pH=7.5 (green) and pH=5.8 (black). NMR spectra were recorded at 0.3 mM RNA concentration, 50 mM K-phosphate buffer, 1 mM MgCl_2_, 90%/10% H_2_O/D_2_O and at 37 °C (a and c) and 25 °C (b).

### 3.4 pK_a_ values of adenosine nucleotides in mismatches differ

We followed the changes in 1D ^1^H NMR spectra during pH titrations for BottomA and BottomB to determine the pK_a_ values of adenine residues (**Fig. 4**). pH values lower than 5.2 (BottomA) and 5.8 (BottomB) were not used in the analysis due to significant broadening of the signals, indicative of pH denaturation of the RNA structures. We followed the proton chemical shifts of the well-defined and resolved signals of A64.H2 (A∙A mismatch) and A58.H8 (G∙A mismatch). While we were not able to directly probe the pK_a_ of A54 (C∙A mismatch) due to signal overlap, we were able to monitor the pH-dependent chemical shift of the neighboring signals, U19.H1 and A53.H2. U19 and A53 form the A-U base pair above the C18∙A54 mismatch and are consequently sensitive to changes in the chemical environment of the C∙A mismatch. As a control we followed the signal of A12.H2 which is present in both BottomA and BottomB and whose chemical shift is not pH-dependent. The observed changes in proton chemical shifts as a function of pH were fitted to **Eq. 1** to obtain the pK_a_ values of the residues. The lowest pK_a_ value was observed for the A∙A mismatch, 5.65 ± 0.05 (A64.H2, **Fig. 4c**). The G∙A mismatch has a pK_a_ value of 6.21 ± 0.05 (A58.H8, **Fig. 4e**). While we could not directly measure the pK_a_ value of A54 (in the C∙A mismatch) due to spectral overlap, we were able to determine pK_a_ values for the neighboring residues U19 (pK_a_ = 6.53 ± 0.05, **Fig. 4g**) and A53 (pK_a_ = 6.66 ± 0.02, **Fig. 4h**). These findings, coupled with the observation that A54 is protonated at pH=7.5, suggests that the pK_a_ of A54 is the highest among the adenines in the base pair mismatches. pH sensitivity is specific for the mismatches in BottomA and BottomB as we observed practically no pH dependence for the A12.H2 signal, which is based paired in the stable region of the stem (**Fig. 4d,f**). Slight changes were observed for A12.H2 in the BottomA RNA due to its proximity to the GAGA tetraloop in comparison to the positioning of A12.H2 in the BottomB RNA.

### 3.5 Thermal stabilization of the structure in C ∙A mismatch but not in others

To evaluate if structure specific differences in pK_a_ values of mismatches are also connected to differences in their thermal stability, we performed UV-melting experiments on BottomA RNA, BottomB RNA, and a series of RNAs harboring single or multiple nucleotide substitutions (**Fig. 6**, **Table 2**). We designed one mutant for the BottomA RNA, BottomA_A8U, that generates a canonical AU base pair at the A∙A mismatch site. For BottomB, we prepared three mutants which either individually stabilized each mismatch (BottomB_G14U, BottomB_C18U) or stabilized both mismatches (BottomB_14/18U). The BottomB RNA, which contains two mismatches, is the least stable at pH=7.5 with a T_m_ value of 56.3 ± 0.1 °C. The stabilization of either one of the mismatches increases the thermal stability of BottomB by almost 10 °C (T_m_ values for BottomB_G14U and BottomB_C18U are 65.3 ± 0.1 °C and 64.1 ± 0.1 °C, respectively). A completely closed stem with both mismatches stabilized in BottomB_14/18U resulted in the highest thermal stabilization (T_m_=73.1 ± 0.2 °C). Similarly, the stabilization of the A∙A mismatch with a U-A base pair in BottomA_A8U shows thermal stabilization of 10 °C (T_m_=71.1 ± 0.3 °C).

**Table 2.** Thermal stability of BottomA and BottomB RNAs.

| Oligonucleotide | $T_m^a$<br>(°C) | | $\Delta T_m$<br>(°C) | $\Delta H^a$<br>(kcal/mol) | |
| --- | --- | --- | --- | --- | --- |
| | pH=7.5 | pH=5.8 | $T_{m(7.5)} - T_{m(5.8)}$<br>(°C) | pH=7.5 | pH=5.8 |
| BottomA | 61.1 ± 0.2 | 59.2 ± 0.1 | 1.9 | -48 ± 1 | -51 ± 8 |
| BottomA_A8U | 71.1 ± 0.3 | 69.8 ± 0.2 | 1.3 | -72 ± 4 | -71 ± 4 |
| BottomB | 56.3 ± 0.1 | 60.6 ± 0.1 | -4.3 | -71 ± 2 | -81 ± 5 |
| BottomB_G14U | 65.3 ± 0.1 | 68.8 ± 0.1 | -3.5 | -90 ± 4 | -91 ± 4 |
| BottomB_C18U | 64.1 ± 0.1 | 64.7 ± 0.1 | -0.6 | -82 ± 3 | -84 ± 4 |
| BottomB_14/18U | 73.1 ± 0.2 | 71.9 ± 0.2 | 1.2 | -93 ± 6 | -92 ± 5 |

<sup>a</sup> $T_m$ and $\Delta H$ values are obtained by fitting UV-melting profiles to two-state model using sloping baselines (Equation 2).

**Figure 6.**
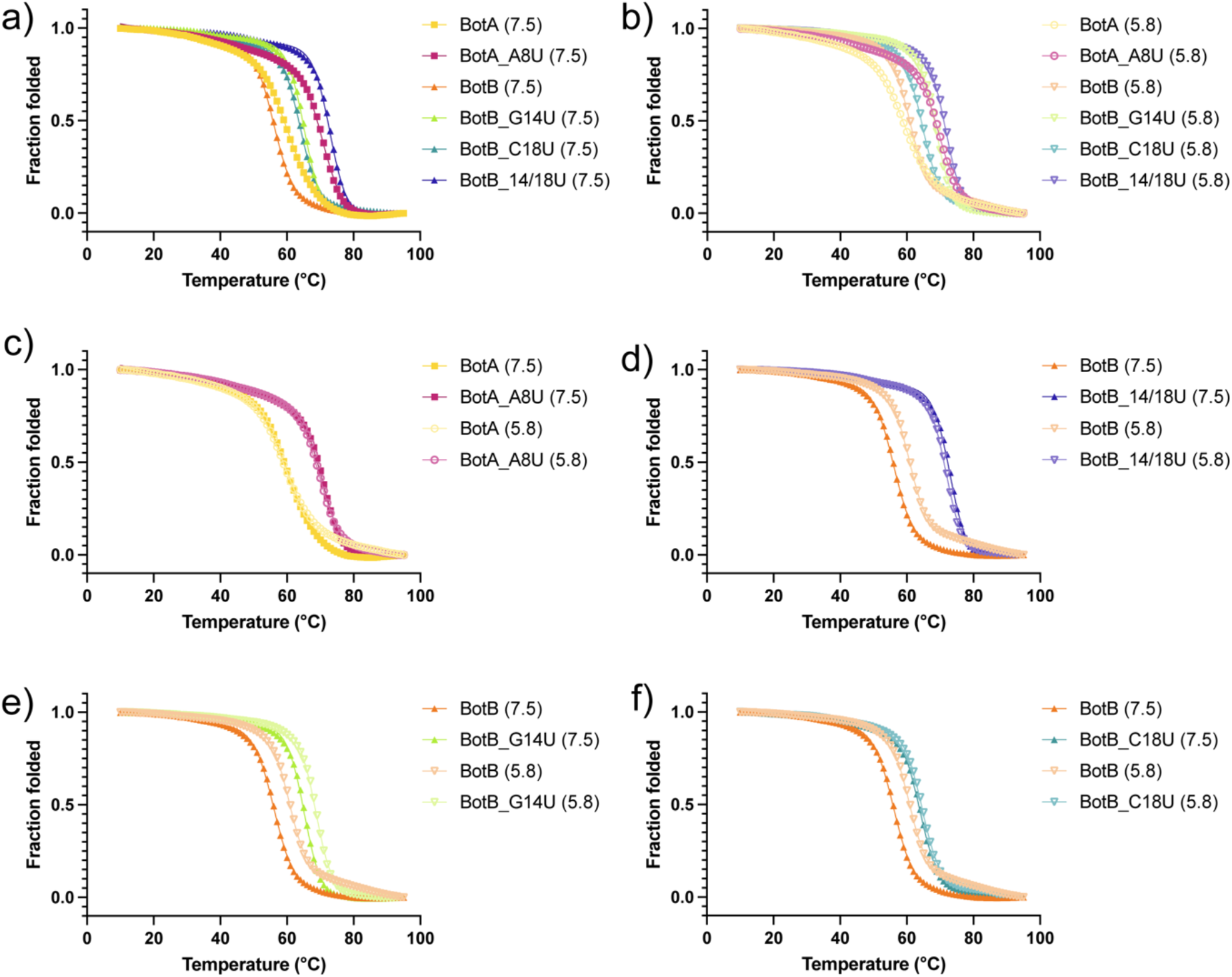
Thermal stability of BottomA and BottomB RNAs. Normalized UV-melting curves (260 nm) of BottomA, Bottom_A8U, BottomB, BottomB_G14U, BottomB_C18U and BottomB_G14C18U RNAs at a) pH=7.5 and b) pH=5.8. Comparison of various mutations at different pH values (c-f). Measurements were performed between 10 and 95 °C on 20 μM RNA in 50 mM K-phosphate buffer (pH=5.8 or pH=7.5). pH values for each sample are indicated in parentheses. Fitted parameters are reported in **Table 2**.

At low pH, we observed slight destabilization of BottomA and BottomA_A8U structures compared to pH 7.5 (ΔT_m_=1.9 °C and 1.3 °C, respectively). Interestingly, at pH 5.8 the thermal stability of BottomB (T_m_=60.6 ± 0.1°C) increased by 4.3 °C relative to that at pH 7.5. This pH-dependent stabilization is connected to the base pair mismatches, since the thermal stability increased by 1.2 °C in the BottomB_14/18U RNA, which has a fully base-paired stem. Interestingly, when the C∙A mismatch is stabilized in BottomB_C18U there is only 0.6 °C increase in thermally stability at lower pH values. The stabilization of the G∙A mismatch in BottomB_G14U resulted in stabilization of the structure by 3.5 °C. These results indicate that the C∙A mismatch is rearranged at lower pH values in such a manner that it contributes to the stabilization of the structure, in which the protonation of the adenine residue A54 is crucial as indicated by NMR data.

## 4. Discussion

Most RNAs contain helical stems with imperfect base pairing, where symmetric (base pair mismatches) and/or asymmetric internal loops disrupt the canonical base pairing. These mismatched regions are thought to mediate important intermolecular interactions, including protein recognition [48–54] and define the global RNA structure [55–58]. Internal loops have also been shown to be important for RNA-mediated processes like ribozyme cleavage [59–61], RNA conformational changes [62–64], and RNA processing [65–68].

G∙A mismatches are the most frequently occurring type of base pair mismatch found in the RNA secondary structure database [69–70] and our analysis revealed that C∙A mismatches are the most common base pair mismatch in the stem of pre-miRNAs. Herein, we investigated the three base pair mismatches, A∙A, G∙A and C∙A, present in the stem of the pre-miRNA-31 **(Fig. 2)**, which is involved in the regulation of a number of physiological and pathological processes. Our NMR-based analysis enabled us to obtain detailed structural information on these base pair mismatches and revealed that each of the three base pair mismatches differ in their structures as well as in their dynamics **(Fig. 3)**. We found the A∙A mismatch to be the most structurally restrained, with both adenine residues oriented within the stem and stacked with neighboring nucleotides. In the case of the G∙A mismatch, we observed a break in sequential NOE connectivity for G14. This is consistent with a structure in which G14 may be flipped outside of the stem. The most flexible and dynamic among the three mismatches is the C∙A mismatch as evident by broadening of the many cross-peaks in the NOESY and ^1^H-^13^C HMQC spectra that define the C∙A mismatch **(Fig. 3)**. Interestingly, similar broadening is also observed in the sequential and intra-residue NOESY as well as ^1^H-^13^C HMQC cross-peaks of U19, A53 and C55 residues which are sequential neighbors of residues A54 and C18 in the C∙A mismatch. The signal broadening observed for the C∙A mismatch and the neighboring residues suggests that the C∙A mismatch is highly dynamic.

The ^1^H-^15^N HSQC spectra clearly show that A54.N1 (^15^N1 labeled A54) is in exchange with its protonated form even at near neutral pH (pH=7.5) **(Fig. 5)**. Indeed, the C∙A mismatch has higher pK_a_ value compared to the A∙A and G∙A mismatches **(Fig. 4)**. The adenosine residues in all three mismatches (A54, A58 and A64) have pK_a_ values above 5.65 ± 0.05, which is higher than the value expected for adenines in single stranded RNAs (pKa ~3.5) [18]. As a control, the pK_a_ values of A54, A58 and A64 residues were also compared to A12, which is positioned in well-defined region of the stem of pre-miRNA-31. In contrast to the three adenine residues found in base pair mismatches, we detected virtually no changes in A12.H2 chemical shifts during the pH titrations.

We can observe a clear correlation between the pK_a_ values of the three mismatches and the differences in their conformations. In the case of pre-miRNA-31, it seems that the higher pK_a_ can be connected to more dynamic behavior of the mismatch in the stem. Additionally, pK_a_ values are influenced by neighboring base pairs and can, in the case of C∙A mismatches, vary between 6.5 and 8.1 due only to effects of nearest neighbors and nearby bulges [71]. Furthermore, a C∙A mismatch in a 20-nucleotide long hairpin with neighboring 5’-GC and 3’-UA base pairs, similar to the mismatch in pre-miRNA-31, has a pK_a_ value of 7.84 and is stabilized by 5.1 °C when the pH value is lowered from 8.79 to 6.13 [71]. These results are in good agreement with the high pK_a_ value of the C∙A mismatch and the 3.5 °C stabilization at low pH in pre-miRNA-31 **(Fig. 6, Table 2)**. Additionally, a small population of the protonated adenine was shown to be reflected in smaller stabilization of the RNA folds at lower pH values [71]. In the case of pre-miRNA-31, the highest stabilization of the structure when lowering the pH was observed for the C∙A mismatch, with even less stabilization for the G∙A mismatch. In the case of the A∙A mismatch, we observed a slight destabilization of the structure.

Protonation of adenine bases is known to play an important role in many biological processes [17, 19, 43–46]. We found that A54 in the C∙A mismatch is protonated in pre-miRNA-31, even at physiological pH and temperature **(Fig. 5)**. This base pair mismatch could impact the stability of the stem and thus could affect the regulation of the Dicer processing. Our studies show that adenine residues can flip in and out of the stem and have different protonation levels, which could have important biological implications. For example, in RNA editing by ADAR, an unpaired adenosine the is “flipped out” of the stem increases editing efficiency [72–73]. Protein-RNA recognition can be modulated by RNA shape and formation of a C∙A^+^ wobble base pair can cause the helical axis of RNA to bend [74]. In the case of pre-miRNA-21, a dynamic equilibrium exists even at physiological conditions where the A^+^∙G mismatch is formed, which enhanced the Dicer processing [17]. It was also strongly implied that the regulation of miRNA biogenesis can be modulated in response to environmental and cellular stimuli [17]. Furthermore, perturbation of the RNA duplex structure by mismatches and wobble base pairs has a negative effect on Drosha/DGCR8 processing [16]. Interestingly, these mismatches and wobble base pairs were not located near cleavage sites but were positioned 10–12 nucleotides downstream of the cleavage sites or 23–26 nt from the basal junction [16]. For TRBP, the protein which recruits pre-miRNAs to Dicer for processing into mature miRNA, it was shown to discriminate between miRNAs based on their secondary structures. TRBP had a strong binding preference for pre-miRNAs, whose stem region had tight base-pairing except for the center region [75–76].

Our studies contribute to our understanding of the chemical and conformational differences between A∙A, G∙A and C∙A in the stem of the pre-miRNA-31. The C∙A mismatch displayed the highest pH dependence as well as a population of protonated adenine residues even at pH=7.5. Lowering the pH value resulted in the stabilization of the C∙A mismatch, had only a slight effect on the G∙A mismatch, and destabilized the A∙A mismatch. Our study suggests that the dynamics of mismatches is connected to their pH sensitivity and might play a role in the regulation of pre-miRNA processing.

## Data deposition

The assigned chemical shifts along with raw NMR data have been deposited in the BMRB. BottomA: 51129, BottomB: 51134.

## Acknowledgements

This work was supported by National Institute of General Medical Sciences of the National Institutes of Health grant R35 GM138279 (to S.C.K.) and the Pew Charitable Trusts Scholars Program (to S.C.K.). Research reported in this publication was supported by the University of Michigan BioNMR Core Facility (U-M BioNMR). U-M BioNMR Core is grateful for support from U-M including the College of Literature, Sciences and Arts, Life Sciences Institute, College of Pharmacy and the Medical School along with the U-M Biosciences Initiative.

