## Supplemental Data for "pH dependence of C•A, G•A and A•A mismatches in the stem of precursor microRNA-31"

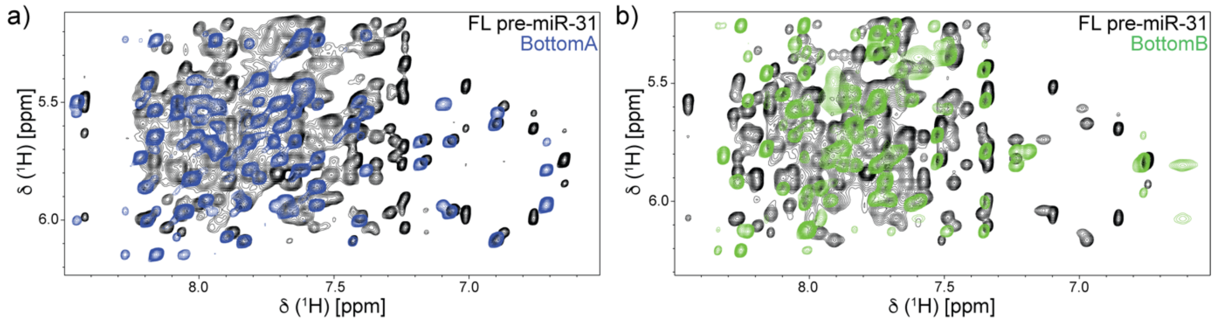


**Figure S1.** Overlay of the aromatic-anomeric region of the $^1$H-$^1$H NOESY spectra for a) BottomA (blue) and pre-miRNA-31 (black) and b) BottomB (green) and pre-miRNA-31 (black). NMR spectra were recorded at 0.4 mM RNA concentration, 50 mM K-phosphate buffer, 1 mM $MgCl_2$, 100% $D_2O$ and at 30 °C (BottomA) and 37 °C (BottomB and pre-miRNA-31).

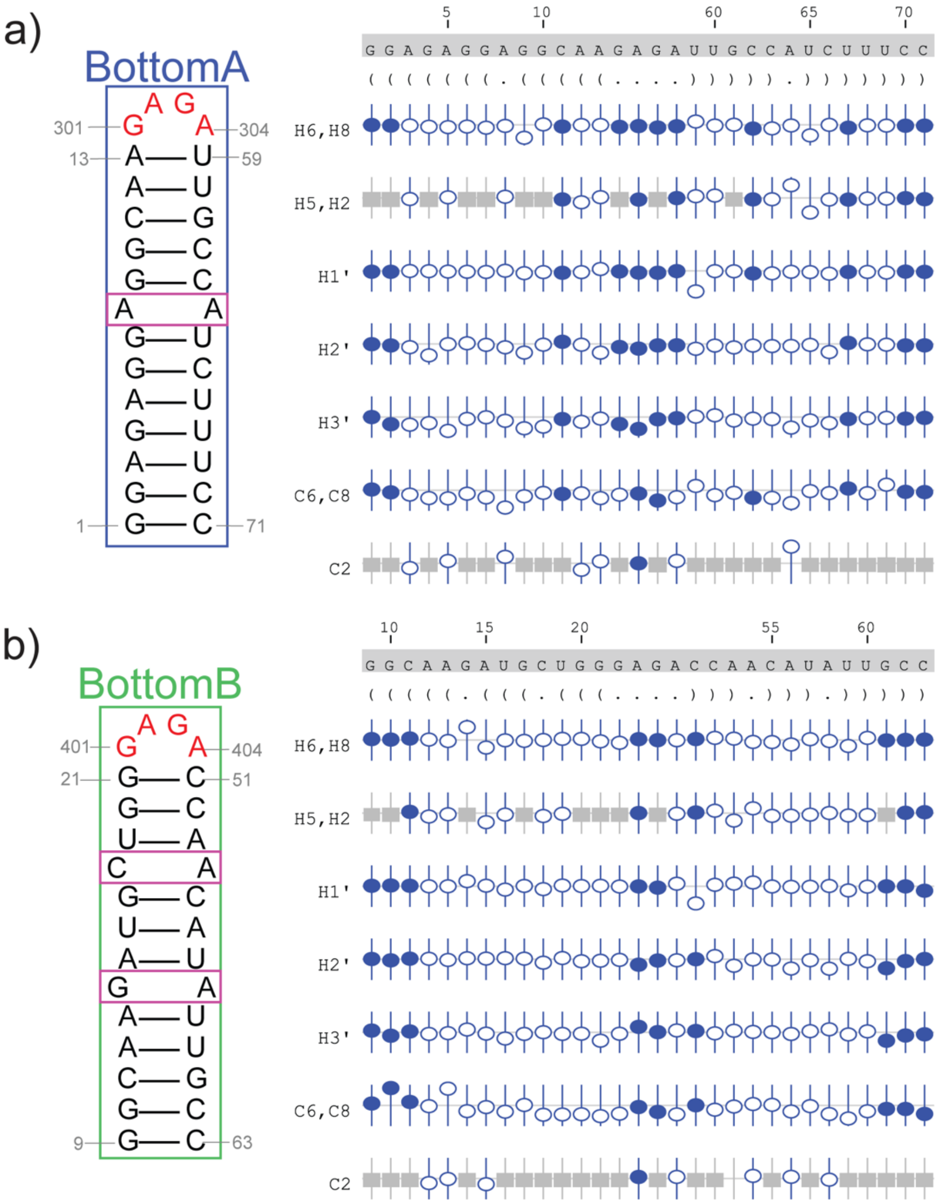


**Figure S2.** Sequence analysis and validation of BottomA and BottomB chemical shift assignments. Secondary structure, nucleotide numbering, and secondary structure in Vienna format for a) BottomA and b) BottomB RNAs. NMRViewJ chemical shift prediction software was used to validate proton (H6/H8, H5/H2, H1´, H2´, H3´) and carbon (C6/C8, C2) assignments. Assigned atoms are represented with blue circles (open and closed), while grey boxes denote atoms that are not present in a given base. Deviation from the predicted chemical shift is shown with the offset from the center. Filled circles indicate that there are chemical shifts for atoms with the same set of attributes in the BMRB. Open circles indicate atoms that have a prediction, but for which no exact matches of the attributes are available in the BMRB.

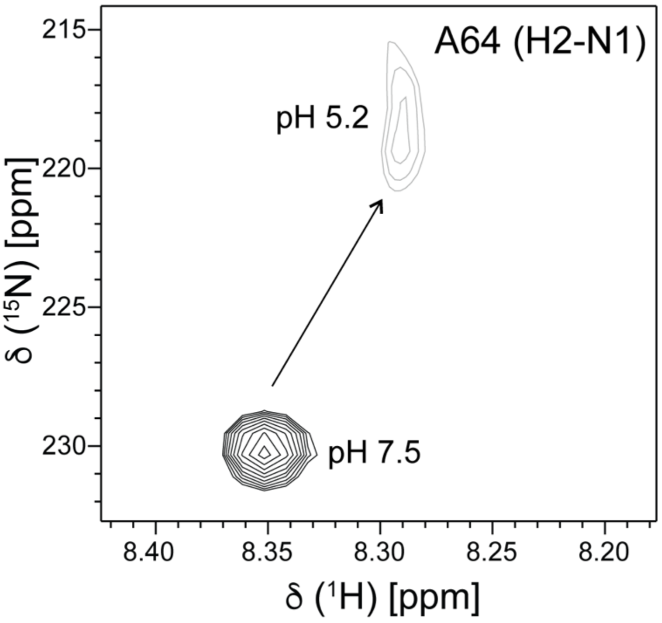


**Figure S3.** H2-N1 cross-peak of 100% $^{15}$N1 labeled A64 in $^{1}$H-$^{15}$N HSQC spectra recorded at pH=7.5 and pH=5.2. NMR spectra were recorded at 0.3 mM RNA concentration, 50 mM K-phosphate buffer, 1 mM $MgCl_2$, 10%/90% $D_2O$/$H_2O$ at 37 °C.

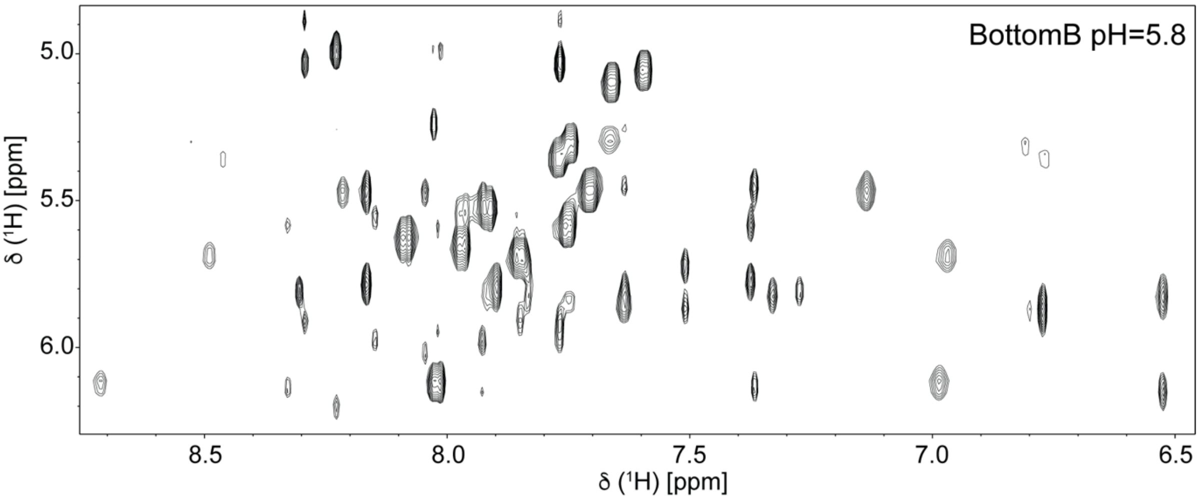


**Figure S4.** Aromatic-anomeric region of BottomB RNA $^1$H-$^1$H NOESY spectrum. NMR spectrum was recorded at 0.3 mM RNA concentration, 50 mM K-phosphate buffer, pH 5.8, 1 mM $MgCl_2$, 10%/90% $D_2O/H_2O$ and at 37 °C.

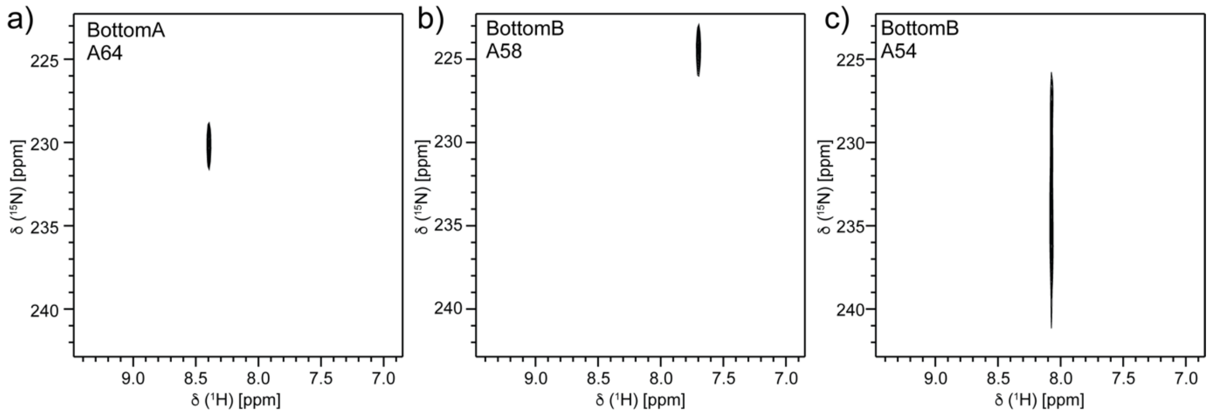


**Figure S5.** H2-N1 cross-peaks of A58 and A54 are much broader compared to A64. H2-N1 cross-peaks of 100% $^{15}$N1 labeled a) A64, b) A58 and c) A54 in $^{1}$H-$^{15}$N HSQC spectra. NMR spectra were recorded at 0.3 mM RNA concentration, 50 mM K-phosphate buffer, pH 7.5, 1 mM $MgCl_2$, 10%/90% $D_2O/H_2O$ and at 37 °C. To obtain NMR spectra of sufficient quality, datasets were recorded with different ns and TD(F1) values as indicated: a) ns=24, TD(F1)=60, b) ns=72, TD(F1)=60, c) ns=272, TD(F1)=76.

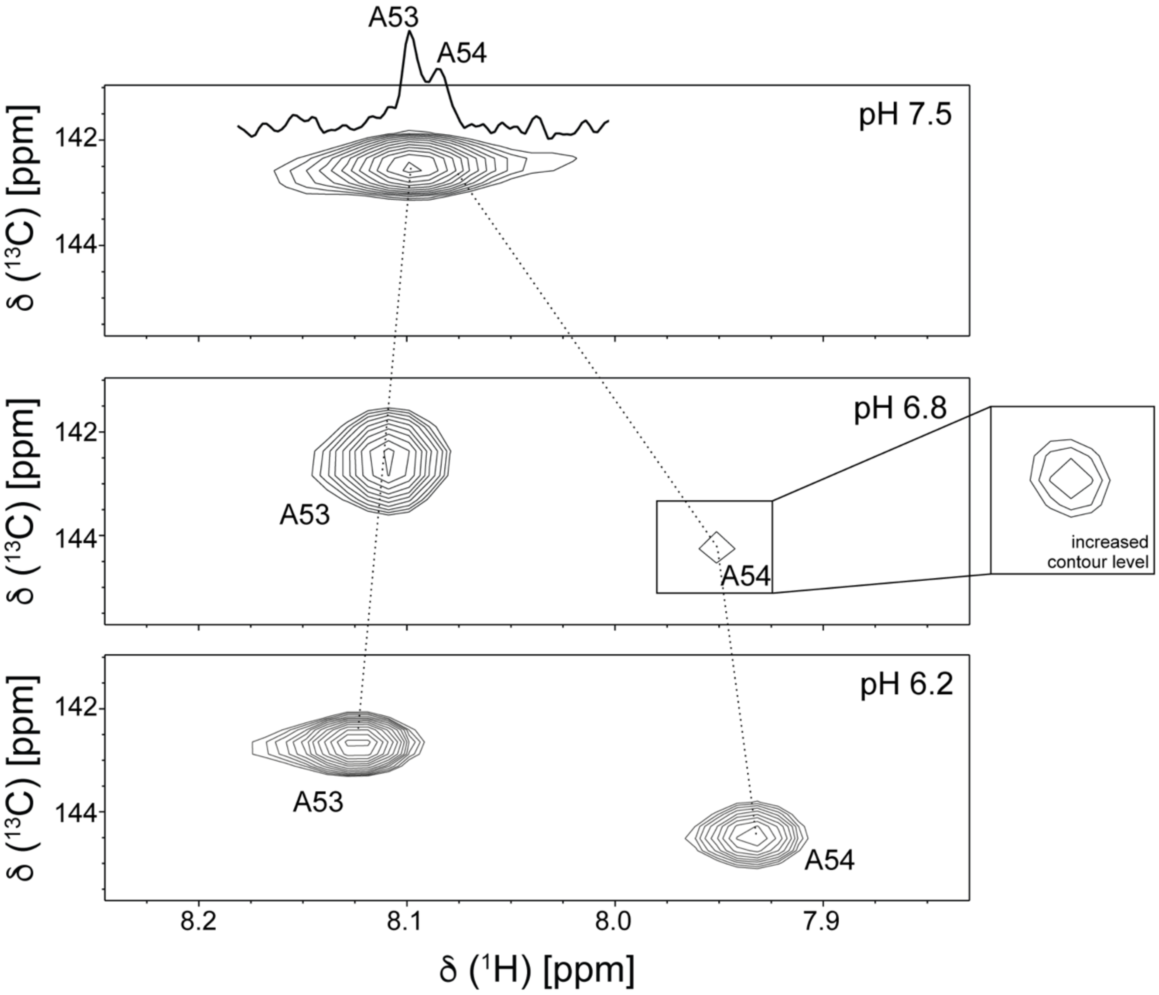


**Figure S6.** H8-C8 correlations in $^{1}H$-$^{13}C$ HSQC spectra of 100% $^{13}C8$ labeled A53 and A54 at different pH values. NMR spectra were recorded at 0.3 mM RNA concentration, 50 mM K-phosphate buffer, 1 mM $MgCl_2$, 10%/90% $D_2O/H_2O$ at 37 °C.
